# Functionally competent CD4^+^ T cells express high levels of T-bet in *Plasmodium chabaudi* infected young mice

**DOI:** 10.1101/2022.02.08.479658

**Authors:** Margaret R. Smith, Komi Gbedande, Corey M. Johnson, Logan A. Campbell, Lyndsay B. Richard, Robert S. Onjiko, Nadia Domingos, Michael M. Opata

## Abstract

The immune system plays an important role in the elimination of *Plasmodium* parasites that cause malaria, which affect children the most worldwide. Immunity to malaria, especially in young children is poorly understood due to the absence of a developmentally-equivalent rodent model to study the pathogenesis of disease. We have developed a mouse model using 15-day old mice (pups) of malaria infection in neonatal mice. Using C57BL/6 pups, we determined that *P. chabaudi* infection decreases the growth rate of young mice compared to controls, and results in 60% mortality, and neurological damage not present in adults, as indicated by a battery of behavioral assays. When all splenic cells were stimulated *in vitro* stimulation, cells from pups proliferated faster than adult cells, but purified CD4 T cells were slower. Upon infection with *Plasmodium* parasites, both adult and pup CD4^+^ T cells were activated and differentiated to an effector T cell (Teff) phenotype; however, pup CD4^+^ Teff were less differentiated than adult Teff. Pup CD4^+^ T cells also produced more IL-2 than cells from adult B6 mice, and TNF-α was increased in parasite-specific BALB/c pup T cells. Interestingly, there were more pup CD4^+^T-bet^hi^ Teff after infection suggestive of increased Th1 commitment, potentially contributing to cerebral symptoms.

## INTRODUCTION

There was an estimated 241,000,000 malaria cases and 427,000 deaths in 2020, with children below the age of five years accounting for 80% of the reported deaths (1). As the disease becomes quickly severe in children, there is high mortality (2). In addition, seizures, coma and other signs of cerebral inflammation during infection predispose children to future disabilities, including poor cognitive development and epilepsy in those that survive the infection (3, 4). While there have been significant advancements in our understanding of malaria pathogenesis, most of these studies have relied on adult animal models, with *Plasmodium chabaudi* being the best described and most physiologically relevant (5). Until now, there has not been a young rodent model of malaria infection. The absence of such a model has limited the growth in knowledge on the disease progression and immune response due to a young immune system.

In establishing a model of malaria in younger mice, we consulted the literature on maturation of mice. A mathematical model has been widely used which relies on the developmental similarities between mice and humans to calculate analogies between mouse age and human age. In this formula, the fusion of growth plates in the scapula is used as a marker of transition from adolescence to adulthood, when humans attain sexual maturity. This makes it possible to determine maturation status by dividing the average sexual maturity for humans in days, by that of mice. Based on this model, it is suggested that 2.60 mouse days are equivalent to one human year (6). Thus, the calculations provided by this model suggest that 14-15 mouse days is developmentally analogous to five human years, a common cutoff used in malaria susceptibility of children, as most cerebral malaria deaths from *P. falciparum* occur before that age. Establishment of such a model will enable the determination of immune development and response to *Plasmodium* infections in young mice. Such information can provide beneficial knowledge for the improved design of malaria treatments and vaccines targeted for children.

Clinical immunity to malaria disease develops more quickly than immunity to the parasites. In malaria, immunity develops gradually as people sustain multiple infections with *P. falciparum*. Clinical immunity reduces disease severity and likelihood of death from the infection (7), while parasite immunity depends on the accumulation of antibodies specific to parasite strains a person is exposed to over time. Levels of malaria-specific antibodies correlate with an increase in age in endemic areas, suggesting an increase due to level of exposure (8). Immunity to parasites requires both B and T cells. The Th1 cytokine IFN-γ can also be measured from peripheral blood mononuclear cells (PBMC) in response to malaria antigens in children presenting with mild malaria (9), and has the potential to facilitate killing of the parasite. While these responses are thought to be beneficial, there are also many regulatory immune processes currently being defined, primarily in T helper cells. For example, malaria patients have unusual Th1-like follicular helper (Tfh) cells which help B cells less than cells with a pure Tfh phenotype, potentially resulting in low antibody levels in malaria-infected children (10). Malaria infection also induces co-production of IFN-γ with the regulatory cytokine IL-10 by CD4^+^ T cells in children with high levels of parasitemia (11). However, IL-21 is also produced by these same cells (12), supporting less than full Th1 differentiation. In some studies, this is accompanied by increases in FoxP3^+^ regulatory CD4^+^ T cells, though Tregs decrease after multiple incidences of malaria (13). These regulatory mechanisms create a balance between pathogen-specific immunity and immune-mediated pathology (14). These immune changes have been studied in *P. chabaudi* infection as well (15, 16). In particular, Th1 differentiation has been shown to be incomplete, with low T-bet, and a pattern of cytokine production described above that is consistent with the persistently stimulated CD4 T cell phenotype called dysfunctional (16). While *P chabaudi* has been well established as a reliable model of pathogenesis and immunity, there should be more investigation of the role of age on immunity and pathogenesis.

Both young mice and children have a lower overall immune response to various stimuli including *Plasmodium* parasites (17–19). However, the mechanisms at play in determining increased severity to disease in young people are not clear, and it is not certain if this is a defect in their ability to develop immunity, or an increase in the propensity to cerebral malaria, which T cells contribute to (20). In this current study, we utilize C57BL/6 mice to describe the immune response to *P. chabaudi* infection in a young rodent model. Such model would be beneficial in understanding the development of immunity to malaria in young children.

## RESULTS

### *Young mice infected with* P. chabaudi *have decreased weight gain*

Children under the age of five can suffer severe malarial disease and are more susceptible to death as a result of *P. falciparum* infection. It has also been proposed that sickness from malaria reduces the growth rate of children, though this is challenging to prove definitively due to the many variables involved with children in endemic areas (21). Therefore, we set out to determine the effect of *Plasmodium* infection on the growth of young mice. To establish the normal growth of pups, we studied the progression of growth in young uninfected mice starting from day 10 after birth for ten weeks. Male and female pups are shown separately but have similar growth patterns **(Fig 1A)**. After 3 weeks (21 days), when we weaned the pups from their mothers, male mice briefly gained weight faster compared to their female littermates. Next, we infected pups at 15 days of age, and adult mice (8 weeks old). Based on a mathematical model by Dutta & Sengupta that determines adulthood and maturity, we approximated that 15-day old pups would developmentally be equivalent to 5.8 human years (6). After infecting the 15-day old pups, we monitored the change in their weight for ten weeks. We observed a significant slowing in weight gain in the infected pups between days 9-21 post-infection (p.i), compared to the uninfected controls **(Fig 1B)**.

**Figure 1:**
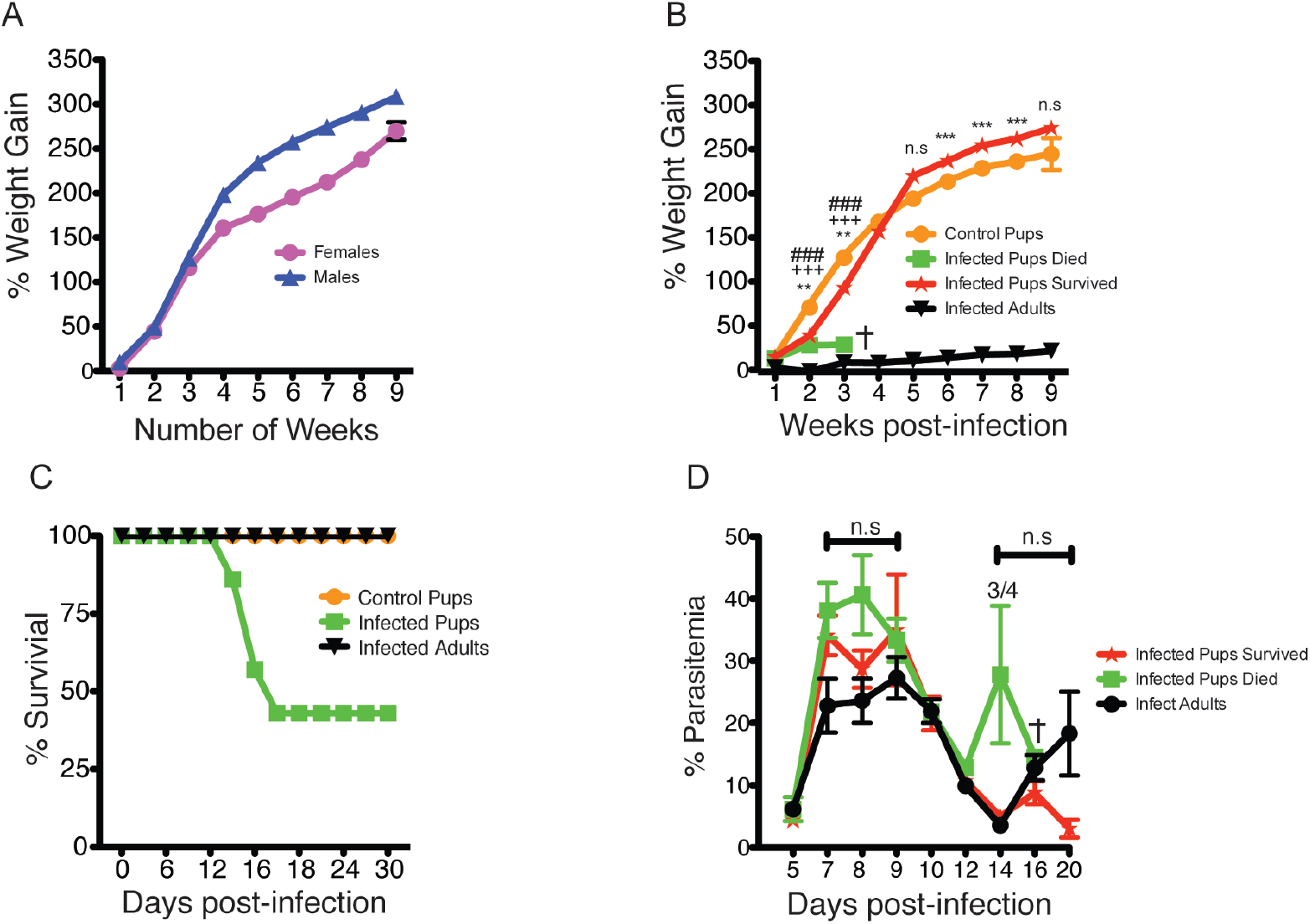
Pups infected with *Plasmodium chabaudi* show a stunted growth and a higher chance of death. (A) Healthy C57BI/6 pups were weighed from day 10 to day 70 post birth to determine weight gain between females and males weight gain. (B) Pups were infected and weights monitored over 9 weeks, data shows the difference in weight of control pups, infected pups that died, infected pups that survived and infected adults. (C) Percent of survival of control pups, adults infected with *Plasmodium chabaudi* and infected pups. (D) Percent parasitemia between pups that survived or died and infected pups from days 5 to 20 post-infection. The data represent three independent experiments with 5-7 pups infected, 3 pups for control and 3 infected adults. The error bars represented the standard error of the mean (SEM). *** p>0.0001, p<0.0022 comparing infected pups that survived to control pups, +++p<0.0030 comparing infected pups that died to infected pups that survived, ###p<0.0002 comparing control pups to infected pups that died using One-Way ANOVA followed by Tukey post-test. Fraction shows number of mice that died out of total mice.

Pups that died strikingly had not gained more than 30% weight by the third week, and pups that lived also had significantly reduced growth. This result is consistent with stunted growth observed in children in malaria endemic areas (21), and suggests that *Plasmodium* infection directly contributes to stunting. Infected adult mice lost weight at the peak of infection, but regained their normal weight after recovery by third week p.i.. Some pups succumbed to the infection, resulting in 60% mortality **(Fig 1C**), and these mice failed to gain weight by the third week of infection. Parasitemia trends showed an increase in pups, but this did not reach significance when compared to parasitemia in adult mice **(Fig 1D)**. Pups that died also had an additional recrudescence at day 14. These data suggest that malaria infection leads to a reduced growth rate of pups during infection.

Malaria can lead to neurological complications and behavior problems in children (22–24). Cerebral malaria can be detected in mice as behavioral changes (25, 26). Therefore, we sought to determine the effects of the infection on the behavior of pups compared to adult mice. Pups (15 days old) or adult mice were infected with a dose of 1×10^6^ *P. chabaudi* infected red blood cells (iRBCs) that is 60% lethal to pups, and performed a animal health and behavior assessment test starting from day 9 p.i.. The SHIRPA, a rigorous and semi-quantitative battery of forty behavioral tests, has revealed significant deficits in animals infected with both *P. chabaudi* (25) and *P. berghei* ANKA-induced experimental cerebral malaria (27). Many scores deteriorated starting near the time of death, and we show only those tests here. In this study, infected pups that died suffered a more severe course of disease which can be measured by significantly decreased scores on day 11 or d14 in several behavioral tests including transfer arousal, tail elevation, body position, spontaneous activity, and pelvic elevation. All of these tests are scored when a mouse is placed in a glass jar with minimal human disturbance. The mice that survived the infection trends towards lower scores in some behavioral parameters, even on day 9, a potential indication that some of these mice also transiently suffered cerebral inflammation. However this was not significantly different from the infected adult controls or uninfected pups **(Fig 2)**.

**Figure 2:**
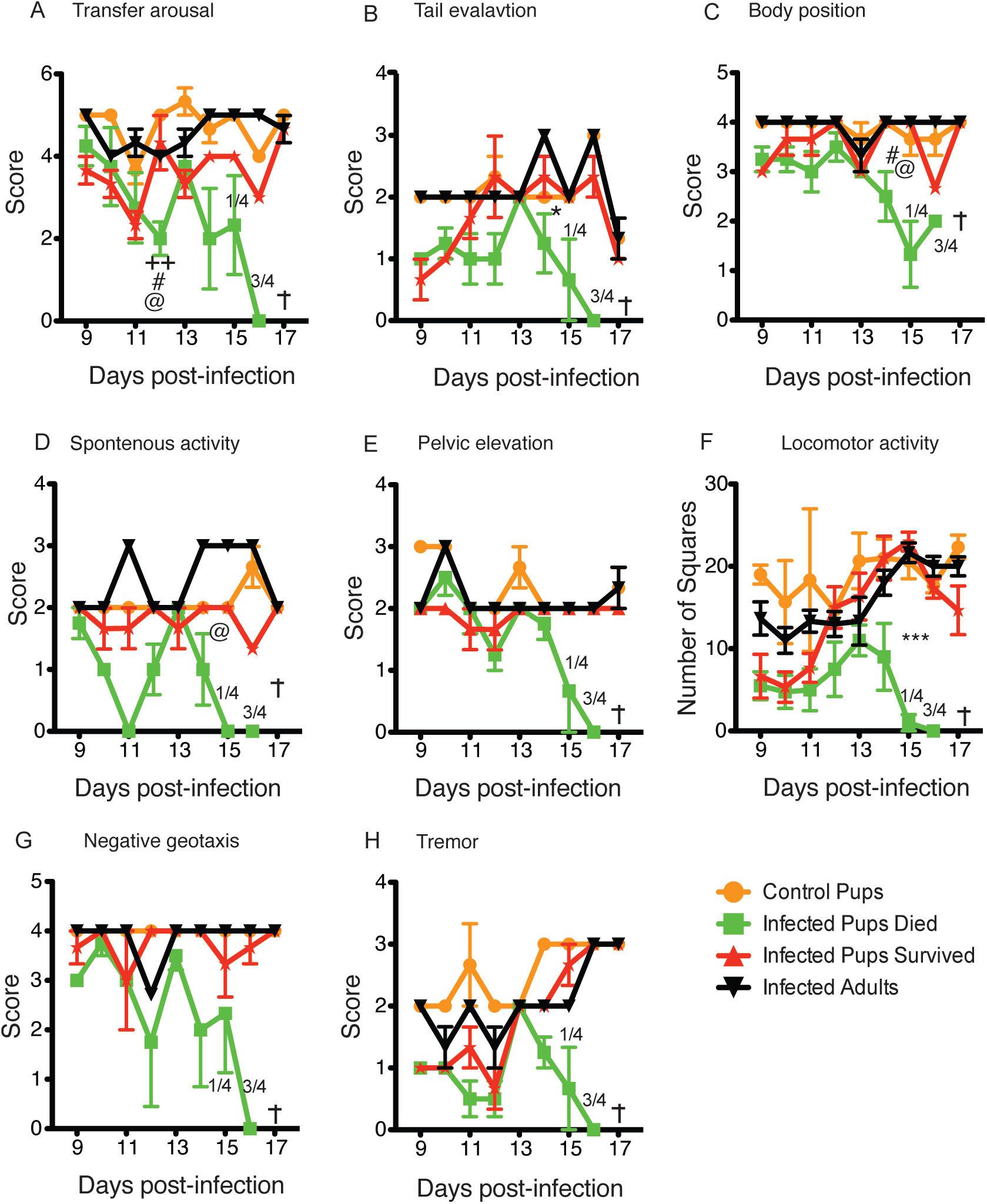
Malaria infected pups have decreased behavior compared to the controls and infected adults. SHIRPA was conducted on in the mice as described in the materials after *Plasmodium chabaudi* infection to understand the natural behavior of infected pups in comparison to adults. Graphs show (A) Transfer arousal to determine the reaction of mice upon quick transfer to a new environment with little to no human contact. (B) Tail elevation (C) body position (D) spontaneous activity and (E) pelvic elevation as the mouse moves around exploring new environment. (F) Locomotor activity when a mouse is placed in an arena and allowed to explore the new environment. (G) Negative geotaxis, when a mouse moves down an inclination, and (H) Tremor when observed in a glass jar. This data represents two independent experiments with 7 infected pups, 3 control pups and 3 infected adults. The error bars represent the standard errors of the mean (SEM). Fraction indicate number of dead mice on the indicated day and + indicate all mice dead on that day. Groups were compared using One-Way ANOVA followed by Tukey post-test with *p>0.05 considered significant. * compares infected pups that survived to control pups, + compares infected pups that died to infected pups that survived, # comparing control pups to infected pups that died, and @ compares infected adults to infected pups that died. Fraction shows number of mice that died out of total mice.

Transfer arousal is a measure of neuropsychiatric state indicative of responsiveness to a new environment (**Fig 2A**) and infected pups were much less responsive than uninfected pups both at day 9 and again by day 15 more significantly with less dramatic changes in adults. Tail elevation (**Fig 2B**) and body position (**Fig 2C**) are measured without human interference as indicators of motor behavioral state. Tail elevation is relatively depressed in infected pups compared to uninfected pups, but not adults at day 9. Both tail elevation and body position scores are dramatically decreased in the pups that die compared to all other groups starting on day 14. Spontaneous activity is a measure of neuropsychiatric state that is significantly decreased only in the pups that are going to die on days 11 and starting on day 14 p.i. (**Fig 2D**). Pelvic elevation and locomotor activity are measures of motor behavior that could be affected by muscle wasting from sickness (**Fig 2E & F**). Negative geotaxis and tremor are behavioral measures that indicate severity of disease **(Fig 2 G & H)**. Some of these changes have not been previously observed in *P. berghei* ANKA or IL-10 deficient mice with *P. chabaudi* infection. Interestingly, it is only decreased from day 14 p.i. in young animals that will die. These data suggest that by the second week of infection, pups show weakened muscular strength and locomotor activity indicators of sickness behavior.

### Pup splenic T cells proliferate faster than adult cells

Since we observed 60% mortality in infected pups, and pup immune system develops slowly, we reasoned that spleen cells from young mice could be defective in response to stimulation. We therefore tested the proliferation capacity of all spleen cells and purified CD4^+^ T cells comparing young and adult mice. We isolated splenocytes and in some cases purified CD4^+^ T cells from either 15-day-old pups or 8-week-old adult mice, and cultured the splenocytes or CD4 T cells on plate bound anti-CD3 and anti-CD28. CFSE dilution was determined by flow cytometry on days 2 and 4 poststimulation to determine the extent of proliferation. By day two of culture, we observed more proliferated spleen cells from the pups, with proliferating cells up to four generations, while adult cells were only in the second generation, with majority of the cells still CFSE positive **(Fig 3A, first panel and graph)**. By day four of culture, all the cells from pup spleens were CFSE negative (CFSE-), while the adult cells were still proliferating in all the four divisions identified **(Fig 3B, second panel and graph)**. On the other hand, purified CD4^+^ T cells from pups had slightly less proliferation than adult CD4^+^ T cells *in vitro*. On day 2 post-stimulation, more than half of the CD4^+^ T cells from the adult mice were CFSE negative, while the pup T cells lagged behind by one generation. In both pup and adult groups, we had cells in all the four divisions **(Figure 3C, left panel)**. The pup CD4^+^ T cells had caught up in proliferation by day 4, although there were still more pup CD4^+^ T cells in the second and third generations than adult cells **(Fig 3D, right panel and graph)**. Taken together, these results suggest that splenocytes from pups proliferate faster, but CD4^+^ T cells may be delayed in their proliferation, even though they are not defective.

**Figure 3:**
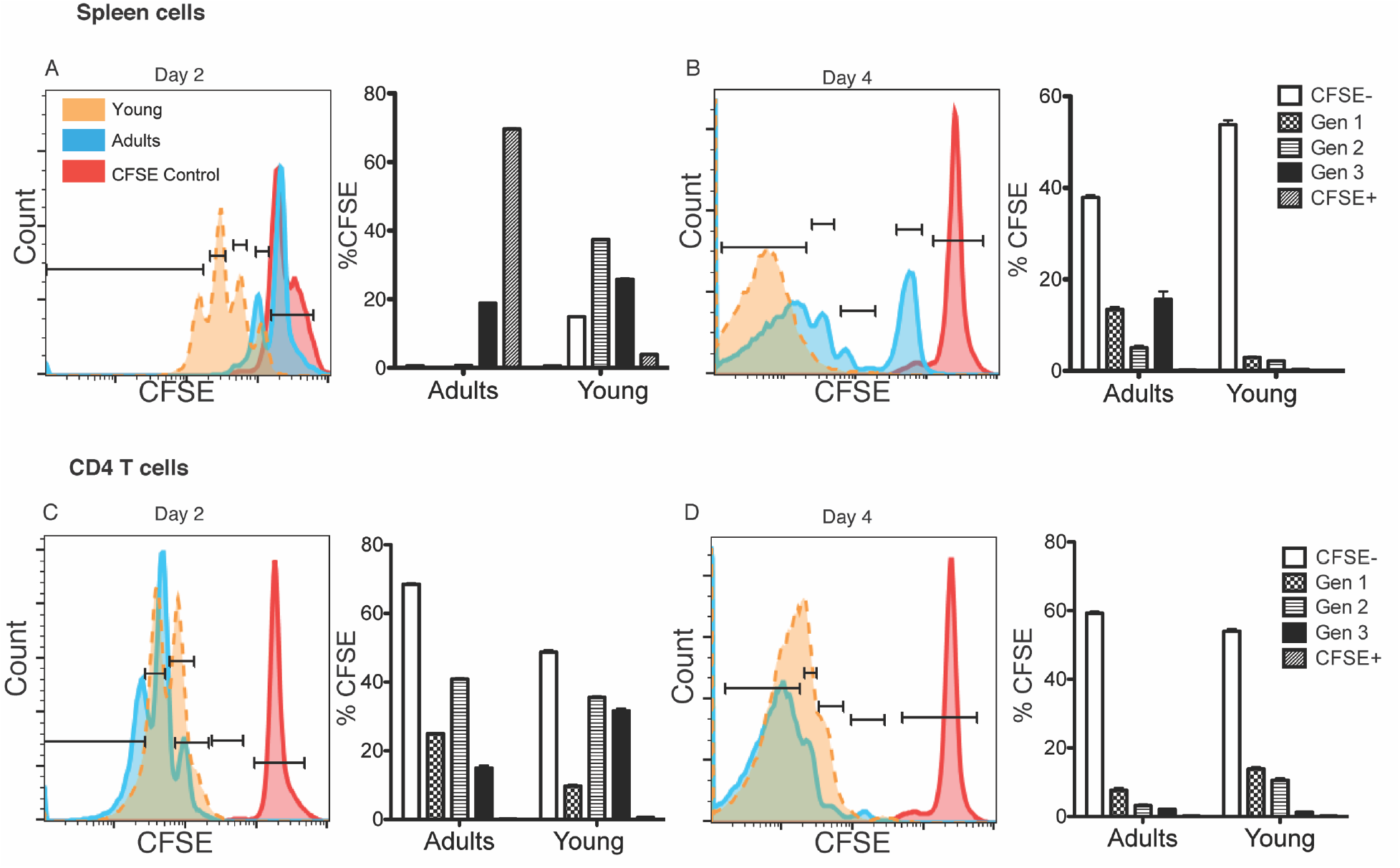
Pup splenocytes proliferate faster than adult cells. All spleen cells or purified CD4^+^ T cells were CFSE-labeled and stimulated *in vitro* using plate bound anti-CD3/CD28. CFSE dilution was measured to determine cell proliferation. (A) Plot and quantified graph showing CFSE dilution in different generations for the spleen cells from adult mice compared to young mice on (A) day 2 and (B) day 4. Plot and quantified graphs showing CFSE dilution in different generation stages of purified CD4^+^ T cells from adult mice compared to young mice on (C) day 2 and (D) day 4. The data is a representative of 5 independent experiments with 3 triplicate wells per experiment. Lines in the plot represent generation stage as quantified in the graphs. The error bars represent the standard errors of the mean (SEM), ***p<0.001, using Student’s t-test.

### Effector CD4^+^ T cells from pups infected with malaria are activated but produce less IL-2 and TNF

Previous studies have suggested that children could be more susceptible to different infections due to the early stage of development of their adaptive immunity (17). To test for differences in CD4^+^ T cell activation and function in response to infection, we infected 8-weeks-old adult or 15-day-old pups with 10^6^ *P. chabaudi* iRBCs, and determined effector T cell activation and numbers on day 8 p.i.. Both pup and adult CD4^+^ T cells were activated as shown by the downregulation of CD127 (IL-7R-) **(Fig 4A-C)** and upregulation of CD44 and CD11a **(Fig 4D-F)**, even though the adult cells were significantly higher compared to the infected pup cells **(Fig 4)**.

**Figure 4:**
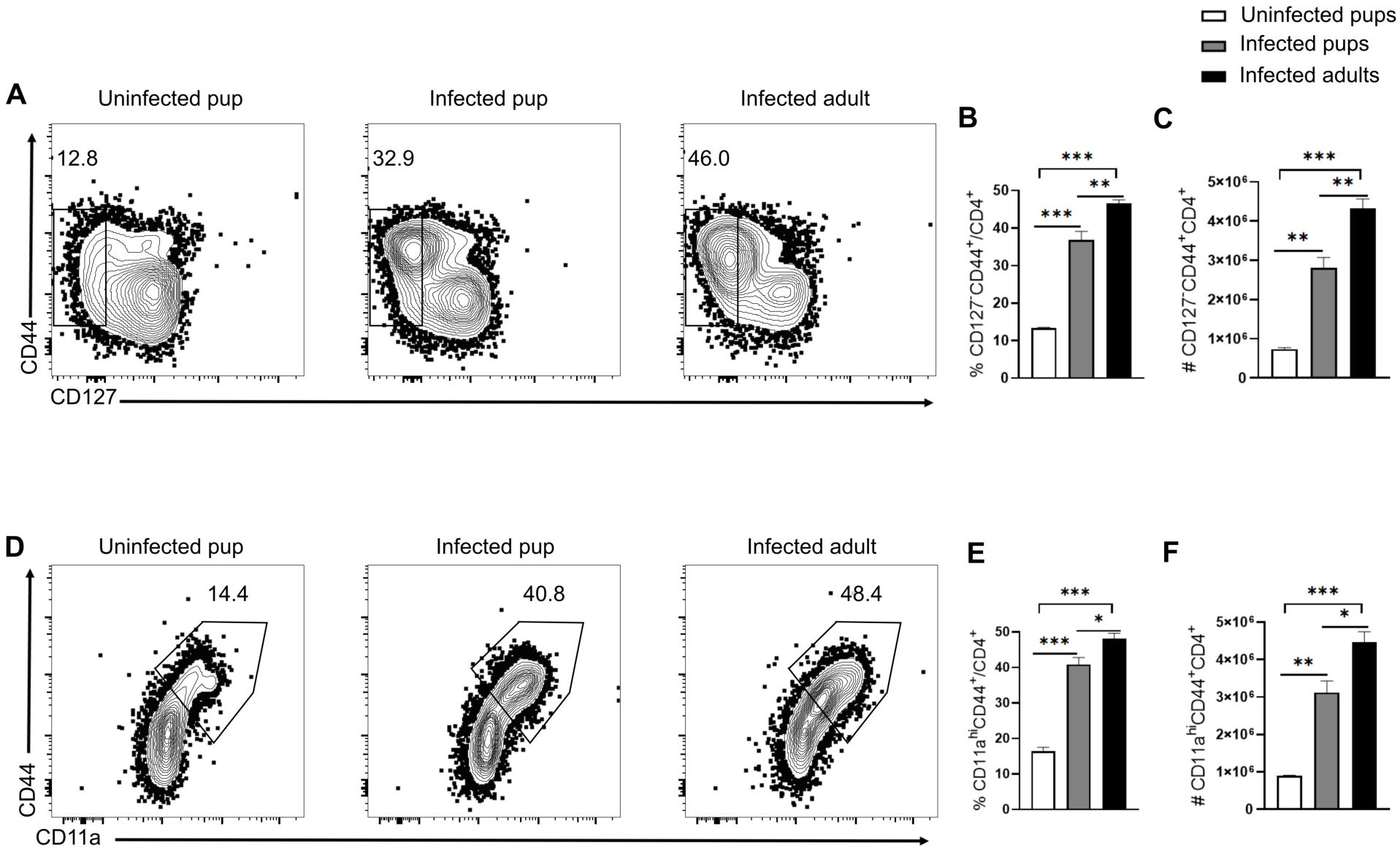
Pup CD4^+^ T cells are activated at a slower rate than adult T cells in response *P. chabaudi* infection. Pups and adults C57BL/6J mice were infected with *P. chabaudi* (10^5^ iRBCs), and CD4^+^ T cells from the spleens were analyzed at day14 post-infection. Uninfected age-matched pups were used as control. (A) Plots and Graphs show (B) percentage and (C) number of Teff (CD127^-^CD44^+^CD4^+^). (D) Graphs and (E) percentage and (F) number of activated T cells (CD11a^hi^CD44^+^CD4^+^). Data are represented as mean ± SEM. One way ANOVA following Tukey’s multiple comparisons test were used to compare between groups, * p <0.05, ** p <0.01, *** p <0.001.

We previously showed a linear differentiation of effector cells into three stages based on CD27 and CD62L expression (28). With lower Teff populations in the pups in Figure 4 above, we wondered if pup cells had any defects in the progression of effector T cell differentiation. While we observed high proportions of Teff^Early^ and Teff^Intermediates^ in the pup cells **(Fig 5A-C)**, the numbers were not significantly different between the infected groups **(Fig. 5E & F)**. The infected adult mice had significantly higher percentages and number of Teff^Late^ **(Fig. 5D & G)**. To measure functional differences in the effector CD4^+^ T cell activation, we tested proliferation-inducing (IL-2) and protective (IL-10, IFN-γ and TNF) cytokine production during *Plasmodium* infection (29–31). Using intracellular cytokine staining, we observed much higher levels of IL-2 in the uninfected pups, which was reduced upon infection **(Fig. 6A)**. Protective cytokines (IFN-γ and IL-10) were increased for both MFI and percentages in the infected groups than the uninfected animals, indicating that cytokine genes were accessible to be transcribed in 6 hours, which is not seen in naïve T cells. Therefore, cytokine positive cells are presumed to be *Plasmodium*-specific and there were no statistical differences between the infected pups and adults **(Fig. 6B & C)**. Interestingly, we observed a significant drop in TNF producing CD4^+^ T cells in all the infected animals **(Fig. 6D)**. These data suggest that pup CD4^+^ T cells are functionally competent and produce similar proportions of protective cytokines to adults’ cells upon infection with *Plasmodium*.

**Figure 5:**
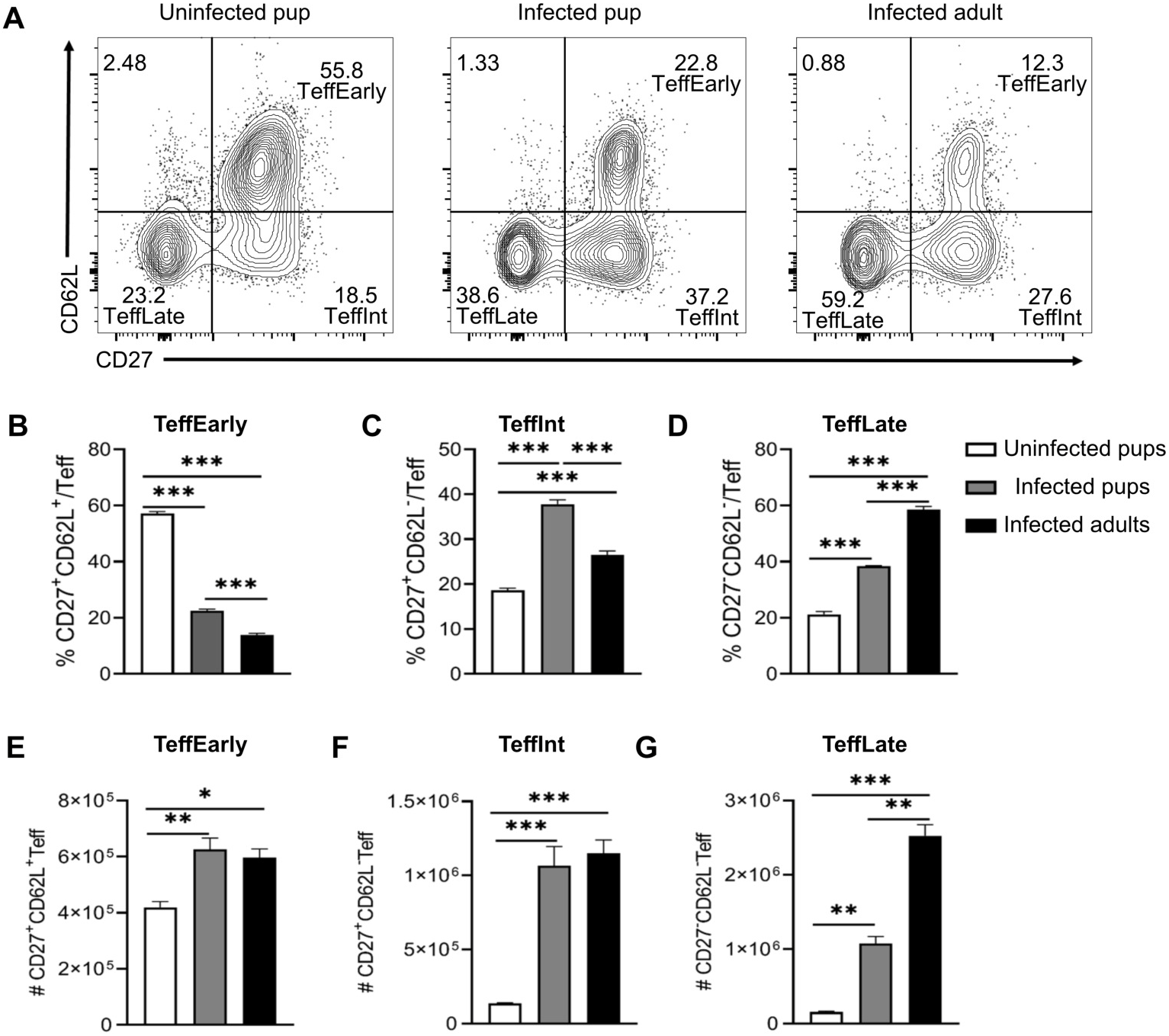
Teff subset differentiation is slower in young mice compared to adult mice after *P. chabaudi* infection. Pups and adults C57BL/6J mice were infected with *P. chabaudi* (10^5^ iRBCs), and splenocytes were analyzed at day14 post-infection. (A) Plots show representative Teff subsets in each group including uninfected pups (left), infected pups (middle) and infected adult mice (right). Graphs below plots show percentages of (A, B, C) or number (D, E, F) of Teff^Early^ (CD62L^+^CD27^+^), Teff^Intermediate^ (CD62L^-^CD27^+^), and Teff^Late^ (CD62L^-^CD27^-^) respectively gated on effector T cells (CD127^-^CD44^+^CD4^+^). Data are represented as mean ± SEM and analyzed using One-Way ANOVA followed by Tukey’s post-test. * p <0.05, ** p <0.01, *** p <0.001.

**Figure 6:**
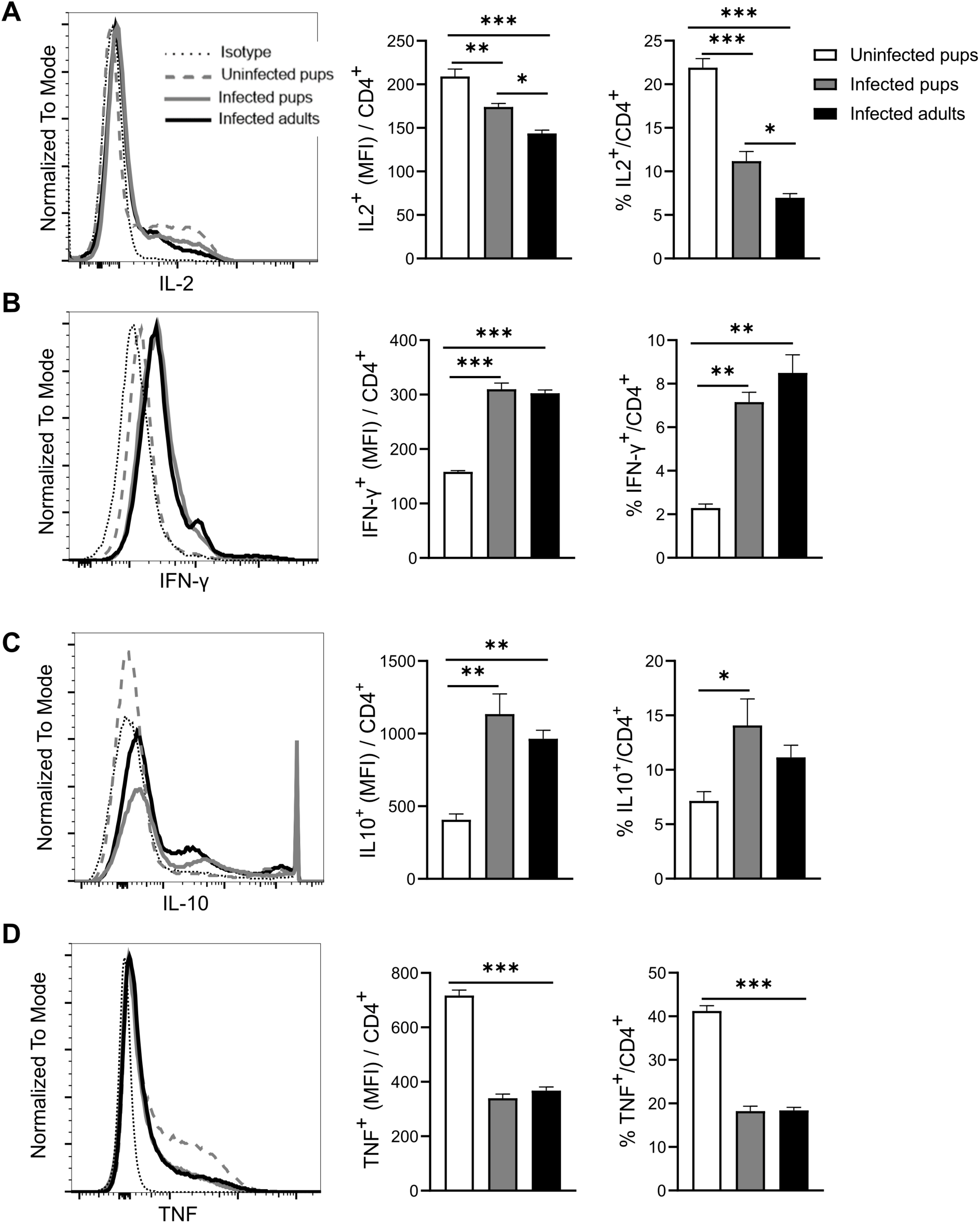
Pup CD4+ T cells secrete more IL-2 but similar level of other protective cytokine with adult mice in response to *P. chabaudi*. Pups and adults C57BL/6J mice were infected with *P. chabaudi* (10^5^ iRBCs), and splenocytes were analyzed at day14 post-infection. Plots show representative histograms of each group and graphs show the MFI and percentage of (A) IL-2 secreting CD4^+^ T cells, (B) IFN-γ secreting CD4^+^ T cells, (C) IL-10 secreting CD4^+^ T cells and (D) TNF-secreting CD4^+^ T cells. Data are represented as mean ± SEM and analyzed using One-Way Anova followed by Tukey’s posttest. * p <0.05, ** p <0.01, *** p <0.001.

To test the hypothesis that there was no difference in the functionality of malaria-specific CD4^+^ T cells between the pups and adult mice, we used adoptive transfer of *Plasmodium*-specific T cells. B5 T cell receptor transgenic (B5 Tg) T cells are specific for *P. chabaudi* Merozoite Surface Protein-1 (MSP-1). Splenocytes from 15-day-old or 8-week-old B5 Tg mice (Thy1.2) were adoptively transferred into congenic adult mice (Thy1.1). Recipients were then infected with 10^5^ iRBCs, one day after cell transfer. The recipient mice were sacrificed on day 8 p.i. to determine *Plasmodium*-specific T cell activation and cytokine secretion. There was no significant difference in the number of recovered cells between the pup and adult Thy1.2 cells **(Fig 7A–C)**. But there were significantly fewer TNF producing CD4^+^ T cells in the recipients of pup splenocytes, while IFN-γ was not significantly different **(Fig 7D & E)**. Taken together, these data show that *Plasmodium*-specific pup T cells are activated, but seem to be significantly defective in TNF secretion.

**Figure 7:**
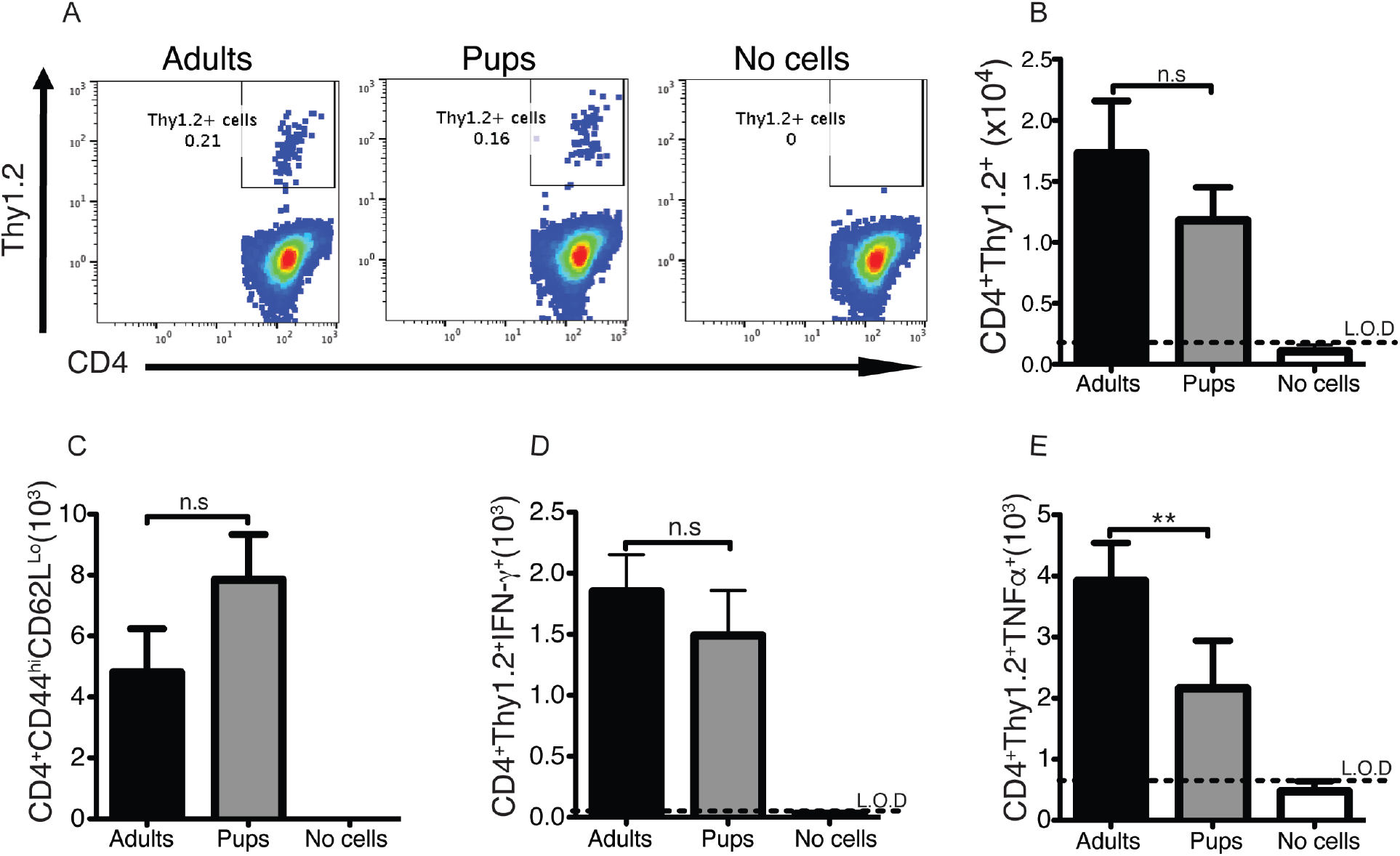
Effector CD4^+^ T cells from adult mice produce more TNFα than pup cells. Splenocytes from adult and pup B5 TCR Tg (Thy1.2) mice were transferred into congenic Thy1.1 mice. The recipient mice were infected with 10^5^ *P. chabaudi* and euthanized on day 8 post-infection to determine CD4^+^ T cell activation and cytokine secretion. **(A)** Plots of recovered Thy1.2^+^ B5 TCR Tg cells. **(B)** Quantification of the recovered CD4^+^Thy1.2^+^ cell numbers. **(C)** Absolute cell numbers of activated effector CD4^+^ Thy1.2+ cells, **(D)** IFN-γ and **(E)** TNFα producing CD4^+^Thy1.2+ *Plasmodium* specific T cells at day 8 post-infection. The data is a representative of 2 independent experiments with 4 recipient mice per group. The error bars represent the standard errors of the mean (SEM).*p>0.05, **p<0.001, n.s - no significance difference, using One-Way ANOVA followed by Tukey’s post-hoc test. “No cells” is a group that did not receive any cells as a control for reconstitution.

### *Pup CD4^+^ T cells express high levels of Tbet after* Plasmodium chabaudi *infection*

Both T helper 1 (Th1) generated by the transcription factor T-bet and forkhead box P3 (Foxp3) regulatory T cells (Treg) are important during blood stage *P. chabaudi* infections (32). These transcription factors facilitate inflammatory responses to eliminate the parasite (T-bet) or modulate immune response to reduce pathology (FoxP3). While T-bet is increased during infection, it does not reach high levels in CD4 T cells in *P. chabaudi* infection, or in human malaria. Foxp3-expressing CD4^+^ Treg are also decreased (33). Pup effector CD4^+^ T cells expressed strikingly higher levels of T-bet after infection, suggesting an actual increase in Th1 cell commitment not seen in adult mice **(Fig. 8A)**. Similar to previous reports, we observed a reduction in Foxp3^+^ CD4 T cells in both the infected and adult mice **(Fig. 8B)**. Taken together with previous studies, these results suggest that despite the reduced number and limited differentiation state of effector CD4^+^ T cells in infected pups, the Teff appear to be highly functional, and strongly polarized to Th1 suggesting a possible mechanism for the increased behavioral symptoms in pups.

**Figure 8:**
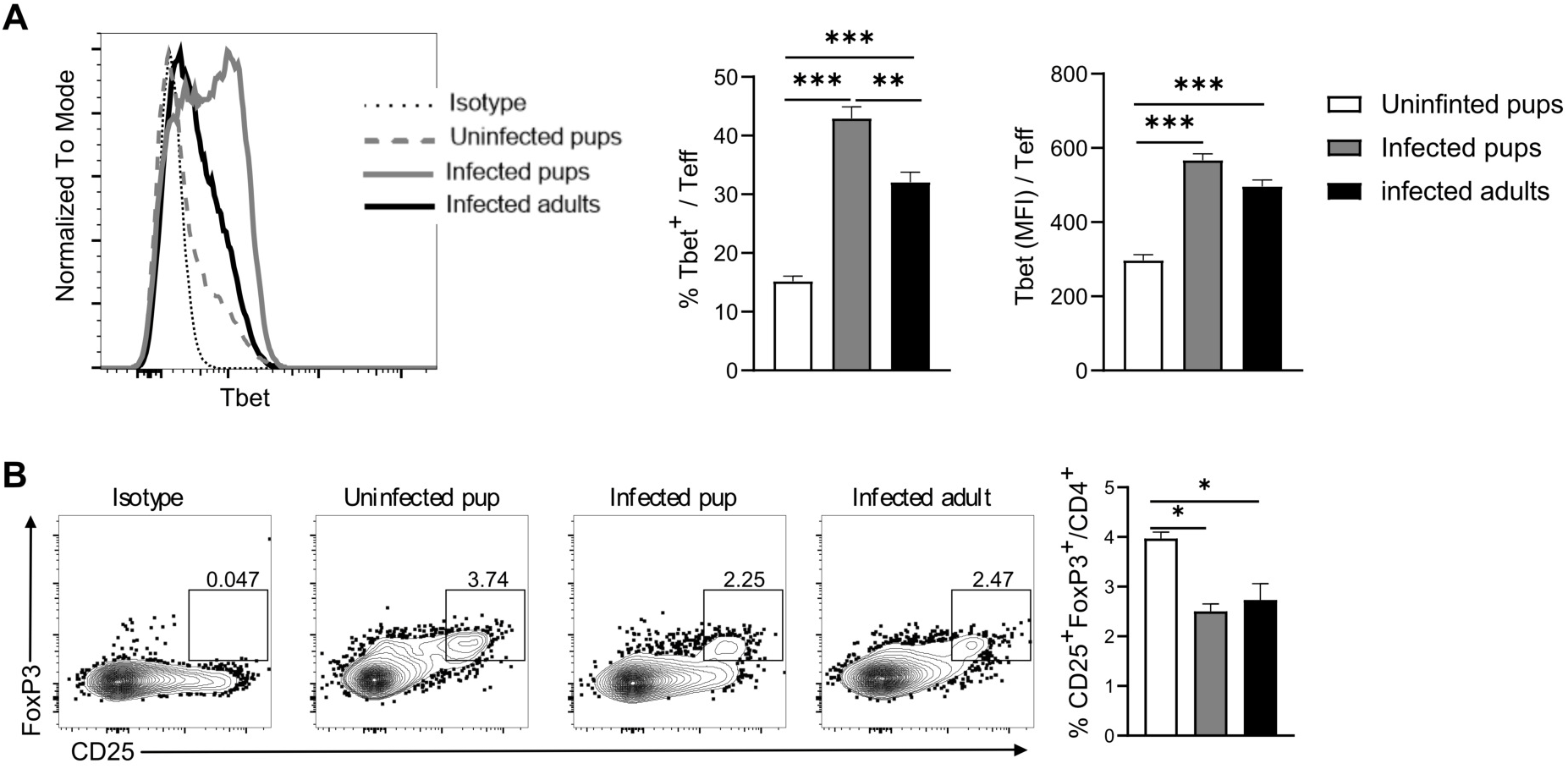
*P. chabaudi* infected pups express high levels of Tbet. Pups and adults C57BL/6J mice were infected with *P. chabaudi* (10^5^ iRBCs), and splenocytes were analyzed at day14 post-infection. (A) Plots show representative histograms of each group and graphs show the percentage and MFI of effector CD4^+^ T cells expressing transcriptional factor T-bet. (B) Plots and graphs show the percentage of regulatory CD4 T cells (FoxP3^+^CD25^+^CD4^+^). Data are represented as mean ± SEM. One way ANOVA following Tukey’s multiple comparisons test were used to compare between groups, * p <0.05, ** p <0.01, *** p <0.001.

## DISCUSSION

Malaria is a significant threat to children in endemic areas. To our knowledge, there is no good established young rodent model for malaria, thus limiting our understanding of this disease in the most vulnerable population. Our current study sought to establish a young rodent model that can be used to aid in the understanding of the immune response to, and pathogenesis of, malaria. While most of the mechanistic knowledge in *Plasmodium* infection to date relies on animal studies (34–37), most of these studies use adult mice. Using our neonatal model, we showed that infected pups had a reduced growth rate during the first 2 weeks of infection. The infection severely reduced weight gain during the first three weeks in 60% of the young mice, which died. The slower growth in the infected pups observed in this study is consistent with a phenomenon seen in human studies, as malaria infection has been reported to increase the risk of stunting in children in many regions including Ethiopia and Lao (21, 38–40), even though these children catch up in growth with their agemates later in adulthood, just as observed in our experiments for pups that recovered **(Fig. 1B)**.

Similar to children who suffer more incidence of severe malaria (41), we observed that more pups than adult mice experience severe disease, as demonstrated by increased mortality and significant changes in behavioral assays **(Figs 2)**. Pups that succumbed to the infection had reduced scores on normal spontaneous activities as well as more complex behaviors such as negative geotaxis, and reduced correcting reflex compared to infected adult mice or pups that survived the infection. These tests are reflective of the changes seen in cerebral malaria in WT mice infected with *P. berghei* ANKA, or IL-10 KO infected with *P. chabaudi* (25–27). These syndromes include extensive cerebral gliosis and vascular events including hemorrhage, edema, and endothelial congestion and coagulation defects throughout the brain, indicating neuropathology (27). Negative geotaxis, or ataxia, and correcting reflex are specifically associated with cerebral malaria and can distinguish the neuro-specific aspects of *P. berghei* infection from more systemic sickness behavior seen.

Understanding the functionality of pup cells can provide important information for vaccine design to malaria in children, the most affected group. Thus, we investigated proliferation, activation, differentiation and cytokine production, which are key functional readouts for protective T cells. Pup CD4^+^ T cells proliferated slower, but they were not completely defective in their ability to proliferate. As expected, effector CD4^+^ T cells were fewer in pups, and differentiated less than adult effector T cells on day 14. The slow differentiation of CD4^+^ T cells could account for the decreased proliferation seen in the stimulated purified T cells from pups. Polyclonal pup T cells produced more IL-2 as measured by percentages and MFI, but there was no difference in IFN-γ and IL-10 production in the effector phase, by both groups. The high IL-2 production in the naive pups may be responsible for the faster proliferation of the splenocytes seen in *the in vitro* stimulation. When we transferred MSP-1 malaria-specific cells into Thy1.1 hosts, there was no difference in the number of recovered (Thy1.2^+^), activated or IFN-γ producing CD4^+^ T cells. However, there were significantly fewer parasite specific TNF producing BALB/c CD4^+^ T cells in the pup donor cells compared to adult TCR Tg T cells, which we did not see in the polyclonal B6 cells.

Higher numbers of regulatory T cells (Tregs) are reported in children from high malaria transmission areas (42). Tregs require transforming growth factor beta (TGFβ) for their development and maintenance, and its presence inhibit inflammatory cytokines important for parasite elimination (43). This could be one reason why children experience severe disease and poor immune response resulting in death from malaria infection. In a recent study, it was shown that Treg cell numbers decline over time in children who are heavily exposed to several malaria infections, which correlates with reduced symptoms (44). Similarly, we observed a decrease in Foxp3^+^ cells in both infected pups and adult mice in our current study supporting an increase in the Th1 to Treg ratio (16).

In humans, it has been reported that infants born to mothers with pregnancy associated malaria during the third trimester of pregnancy had a significantly increased risk of infection during the first year of life (45). Contrary to the observation in humans, a recent study by Duffy’s group showed that pups born of infected pregnant mothers and infected after weaning as adults (weeks 6-8 after birth), are resistant to infection and are protected from weight loss (46). Using our newly developed model, it would be interesting to infect the pups exposed to malaria during pregnancy at a younger age of 15 days to see if the parasitemia or frequency of cerebral symptoms noted here is affected. Such investigations are likely to help in identification of procedures that would induce protection in neonates, and could further expand our knowledge of *Plasmodium* infection, allowing application of science to reduce death rates in vulnerable populations.

In conclusion, our data suggest that in response to *Plasmodium* infection, there are fewer effector CD4^+^ T cells generated in pups, and that they have a less differentiated phenotype by day 14, however, they have significantly higher T-bet expression which may facilitate production of protective cytokines to the same level as adult cells. Several cytokines, including IFN-γ were produced at largely similar levels using intracellular cytokine staining, therefore T-bet levels may indicate a more highly committed Th1 phenotype. Interestingly, animals deficient in IL-12Rβ2, which is generally required for full Th1 commitment and high levels of T-bet, are less susceptible to cerebral malaria from *P. berghei ANKA* (47), as are T-bet deficient mice (48). Interestingly, T-bet does not appear to be required for control of parasitemia (48). This increase in T-bet is strikingly different to the changes in Th1 response in neonatal mice to Listeria, an infection that causes strong Th1 commitment, but where T-bet and IFN-γ were down in pup splenocytes by real time PCR (49). In polyclonal CD4^+^ T cells in B6 mice, IL-2 was higher in pups, but *Plasmodium*-specific T cells produced less TNF.

While further work needs to be done to establish the role of CD4 and CD8 T cells in the brain of infected pups, both of these cells make IFN-γ that is essential to experimental cerebral malaria in adult mice, and both T-bet and IL-12Rβ2, a marker of Th1 commitment are required for experimental cerebral malaria. We speculate that the less differentiated Teff phenotype in our study, suggests an interesting pup T cell bias towards multipotency, another interesting finding to follow up using this new model. Thus, our findings provide a good young rodent model to help in understanding the pathogenesis of malaria in the population that is most affected with the disease as 80% of deaths reported for malaria occur in children (1). Such knowledge on malaria may be beneficial to other chronic parasitic infections like *Leishmania*, tuberculosis among others. This could improve vaccine design targeting protective cells in different diseases thus reduce neonatal mortality.

## Acknowledgements

The authors would like to thank all members of the Opata lab specifically Jennifer Pilotos, Hugo Santos and Nadia Domingo for their technical assistance with various experiments. We’d like to thank Monique Eckerd and Scott Rhymes for their help with animal husbandry, Drs. Seals and Ahmed for access to the flow cytometer in their lab and Drs. Mark Robinson and Robin Stephens for reading the manuscript for clarity. These studies were funded by the Department of Biology, Office of Student Research and College of Arts and Sciences at Appalachian State University; and Supported by the James W. McLaughlin Fellowship Fund (KG). The funders had no role in study design, data collection and interpretation, or the decision to submit the work for publication.

## MATERIALS AND METHODS

### Mice and parasites

C57BL/6 mice were maintained in a breeding facility at Appalachian State University Animal Facility Lab, or purchased from Charles River Laboratories (Wilmington, MA). Young mice were generated by breeding. Young mice were 14 to 16 days old of age, while adult mice were 8 weeks when infected with *P. chabaudi* (*AS*) for all experiments. All experiments were carried out in accordance with the protocols approved by the UTMB Institutional Animal Care and Use Committee and The Appalachian State University Institutional Biosafety Committees.

### SHIRPA

SmithKline Beecham, Harwell, Imperial College, Royal London Hospital, phenotype assessment (SHIRPA) was conducted to characterize the behavioral changes in pups during three stages with a series of individual tests that provide quantitative data about the pup’s individual performance (25, 50). SHIRPA mimics the general and neurological examination in humans, and measures Reflexes and Sensory Function, Neuropsychiatric Function, Muscle and Lower Motor Neuron Function, Spinocerebellar function, Autonomic Function, as well as Muscle Tone and Strength. When a pup showed a poor performance in a locomotor test early in infection this reflected a muscular weakness due to a dysfunction in the central nervous system, not muscular wasting seen later. The tests were run in a simple manner that did not change the course of infection, and we allowed them 30 minutes rest before another test.

### SHIRPA tests to assess Strength and Sickness

The mouse is transferred quickly into a new environment with little to no human contact in order to observe the immediate reaction. For transfer arousal a score of 5 showed that the mouse did not freeze when in a new environment while a lower score indicates a longer pause before moving. For tail and pelvic elevations, the mouse is observed as it explores its new environment during its forward motion. A score of 2 indicate that the tail is elevated, a 1 indicating it is extended horizontal, and a score of 0 shows that the tail is being dragged. If the pelvic region was more than 3mm in elevation the score of 3 was assigned, 2 represented a normal elevation of 3mm, 1 showed that the pelvic elevation was barely touching the floor and 0 was the pelvis was flattened on the ground. Mice were in the beaker for 5 minutes before body position was recorded. A score of 4 indicated the mouse was standing on their hind legs, a 3 meant the mouse was sitting, a 2 was lying prone and when the mouse was lying on its side it was a score of 1. When the mouse placed in the beaker it is observed for spontaneous activity. A score of 3 shows that the mouse had a rapid and darting movement, a score of 2 showed vigorous grooming and moderate movement, 1 representing casual grooming or slow movement. When a mouse was resting or showed no movement it was given a score of 0. As the mouse was exploring the new environment the pelvic elevation was observed.

### SHIRPA tests to test Neurological Deficits

The mouse was placed in an open arena and allowed to explore the new environment. The arena had equal length and width in squares. For locomotor activity, the number of squares all four paws entered was counted over 30 minutes. For tremor, the mouse was observed when placed in a glass beaker. A score of 2 shows no tremor, a 1 represents a mild tremor and a 0 showed that an important tremor was observed. For negative geotaxis, a mouse was placed on a horizontal cage top and when the mouse moved in one direction the top was raised vertically so that the animal was facing downwards. A stopwatch was set for 30 seconds, and the mouse was observed. A score of 4 showed the mouse turned around and climbed up the grid, a 3 represents a mouse that turned around but froze, when a mouse moved but did not turn around it was given a score of 2, 1 was given when the mouse did not move for 30 seconds and 0 represents when the mouse fell off grid. For touch escape, a mouse was stroked by a finger while exploring its new environment. If the mouse vigorously escapes the finger stroke it was given a score of 3, if the mouse had a rapid response to a light stroke it was given a score of 2, a score of 1 indicated that a firm stroke was required to get an escape response. For visual placing, the mouse is held by its tail and lowered to the cage top the extension of the forelimbs by the animal was observed. A score of 4 showed that the mouse had early vigorous extension around 25mm above the cage top, 3 represents an extension of forelimbs before vibrissa contact around 18mm, 2 shows the extension of forelimbs occurred upon vibrissa contact, 1 means it occurred upon nose contact and 0 shows that there was no response.

### Monitoring Weight

Adult and young mice were infected with 10^6^ *P. chabaudi* parasites. Mice were weighed daily on a balance. Percent weight change was calculated using the original weight on the day of infection [(Weight today – weight d0)/Weight d0] x 100%

### Flow cytometry

Single-cell suspensions were obtained from the spleens in complete Iscove’s media (5% FCS, and 3mM HEPES). Cell pellet was resuspended and incubated in red blood cell lysis buffer to eliminate red blood cells. Cells were then washed in phosphate-buffered saline supplemented with 2% fetal bovine serum, and 0.01% sodium azide, then surface stained with anti-CD4-fluorescein isothiocyanate (FITC), anti-CD44-phycoerythrin (PE) and anti-CD62L-phycoerythrin/Cyanine 7 (PE/Cy7). Data was collected using the FC500 flow cytometer (Beckman Coulter or Forttesa BD) and data analyzed using the FlowJo software (BD/TreeStar, Ashland, OR).

### Intracellular Cytokine Staining

Single cell suspensions were enumerated 5×10^6^ cells stimulated with 1X of cell stimulation cocktail (Tonbo Biosciences, San Diego CA) for 5 hours. The cells were cultured for 5 hours at 37°C, 5% CO_2_ in the presence of GolgiPlug (BD Biosciences) for intracellular cytokine staining. Upon harvest, cells were surface stained with CD4 eflour 450, CD25 PE/Cy7, CD44 APC-efluor 780, CD127-PE/Cy5, and incubated for 30 minutes. Samples were washed with FACS buffer (PBS with 2% FBS and 0.01% Sodium Azide). Cells were fixed by adding 2% PFA in PBS followed by a 30 minute incubation. The cells were then incubated in 1X Perm/Wash buffer (Tonbo Biosciences) for 30 minutes to permeabilize the cell membrane. After permeabilization, cells were incubated with anti CD16/32 Fcblock (BioXcell, Lebanon, NH) for 20 minutes, followed by addition of anti-IFN-γ FITC, IL-2-PE, IL-10-BV650, TNFα-PE/Cy7, for cytokine intracellular staining and T-bet-PerCp-Cy5.5, FoxP3-FITC using transcription factor staining buffer (ebiosciences) for transcription factor staining. Then cells were incubated for a further 40 minutes in the refrigerator (8-12°C). Cells were then washed 3 times with perm/wash buffer, and samples were analyzed by the Fortessa or FC500 flow cytometer. Data was analyzed using FlowJo.

### Adoptive transfer

Spleens were collected from or 8-week-old B5 TCR Tg mice (if infected then day 14 p.i.) and mashed into single cell suspension using a mesh-screen. Red blood cells were lysed using 1X lysis buffer (Tonbo Biosciences). Cells were then counted and 1×10^7^ splenocytes were transferred intraperitonially into Thy1.1 congenic mice. The recipient mice were infected with 10^5^ *P. chabaudi* and sacrificed on day 8 to determine CD4^+^ T cell activation and cytokine production.

### In vitro culture

Corning 24 Well plates were coated with 1μg/ml of anti-CD3 and CD28 then incubated overnight at 4°C. CD4 T cells were purified from the spleen cells using CD4 selection kit from STEMCELL Biotechnologies (Vancouver, Canada). The purified CD4 T cells or unpurified splenocytes were labelled with 1μM of CFSE (Tonbo Biosciences, CA), by incubating in a water bath at 37°C for 10 minutes. Labelled cells were cultured in the pre-coated 24 well plates at a concentration of 1×10^6^ cells/well. The cells were harvested on days 2 and 4 and enumerated then the proportion of proliferated cells were determined based on CFSE dilution on an FC500 flow cytometer.

### Statistical analysis

Data collected after FlowJo analysis was quantified in Microsoft excel and statistical analyses performed using GraphPad Prism (TreeStar, Where) version 5.0 for Mac. Statistics was determined based on standard error of the mean. P values were calculated using one-way ANOVA followed for Tukey posthoc test where necessary due to non-normality of the data. In some experiments, unpaired Student’s t-test was used and p ≤ 0.05 was considered significant.

## List of Abbreviations

ACT: artemisinin combination therapy
IFN-γ: interferon gamma
TNFα: tumor necrosis factor
IL-2: Interleukin 2
IL-10: Interleukin 10
TGFβ: transforming growth factor beta
Pups: young mice
ITN: insecticide-treated nets
PBMC: peripheral blood mononuclear cells
SHIRPA: SmithKline Beecham, Harwell, Imperial College, Royal London Hospital, phenotype assessment

